# Covid19Risk.ai: An open source repository and online calculator of prediction models for early diagnosis and prognosis of Covid-19

**DOI:** 10.1101/2021.01.05.425384

**Authors:** Iva Halilaj, Avishek Chatterjee, Yvonka van Wijk, Guangyao Wu, Brice van Eeckhout, Cary Oberije, Philippe Lambin

## Abstract

**Objective:** The current pandemic has led to a proliferation of predictive models being developed to address various aspects of COVID-19 patient care. We aimed to develop an online platform that would serve as an open source repository for a curated subset of such models, and provide a simple interface for included models to allow for online calculation. This platform would support doctors during decision-making regarding diagnoses, prognoses, and follow-up of COVID-19 patients, expediting the models’ transition from research to clinical practice.

**Methods:** In this proof-of-principle study, we performed a literature search in PubMed and WHO database to find suitable models for implementation on our platform. All selected models were publicly available (peer reviewed publications or open source repository) and had been validated (TRIPOD type 3 or 2b). We created a method for obtaining the regression coefficients if only the nomogram was available in the original publication. All predictive models were transcribed on a practical graphical user interface using PHP 8.0.0, and published online together with supporting documentation and links to the associated articles.

**Results:** The open source website https://covid19risk.ai/ currently incorporates nine models from six different research groups, evaluated on datasets from different countries. The website will continue to be populated with other models related to COVID-19 prediction as these become available. This dynamic platform allows COVID-19 researchers to contact us to have their model curated and included on our website, thereby increasing the reach and real-world impact of their work.

**Conclusion:** We have successfully demonstrated in this proof-of-principle study that our website provides an inclusive platform for predictive models related to COVID-19. It enables doctors to supplement their judgment with patient-specific predictions from externally-validated models in a user-friendly format. Additionally, this platform supports researchers in showcasing their work, which will increase the visibility and use of their models.

## Introduction

The recent COVID-19 pandemic, at its start, emphasized several key unmet needs in terms of patient stratification using quantifiable metrics [1]. These include (a) identifying, in the uninfected population, at-risk persons who should be subjected to stricter restrictions than the general population [2], and (b) in the infected population, improving the detection of high-risk patients by utilizing all available patient data (e.g., clinical, laboratory, genetic, and radiological features) so as to improve quality of care and use of hospital resources [3][4]. Now, with several vaccines emerging, there is another compelling reason for identifying those who are most at risk and should therefore receive the vaccines first [5–7].

Ideally, one should address the above needs using quantitative tools that (a) help people at home decide (in consultation with their doctor) whether their health status warrants being self-quarantined, and whether their symptoms (if present) indicate the need for visiting the hospital, and (b) help doctors during triage decide if a patient should be sent home, hospitalized in a ward, or admitted to intensive care [8]. Quantifying these probabilities can be done by using predictive machine learning models.

Currently, COVID-19 publications regarding such models are booming. There are numerous studies being published, from multiple countries and all using different inclusion criteria and outcome measures [4]. This heavily complicates the selection of the optimal model for a specific patient [9]. In addition, the quality of the research is sometimes suboptimal, as a recent review paper has shown [4].

We, as researchers working on COVID-19 models, saw an urgent need for a web-based platform that would serve as an open source repository for validated models. Such a platform would allow the user to have a quick overview of the strengths and weaknesses of the curated models that passed our quality checks. The platform would also allow the user to calculate the output of such models by simply providing the inputs in a user-friendly format, rather than creating their own implementation or conducting their own search to find a suitable implementation.

Our aim for this platform is to include validated prediction models (TRIPOD type 2b and 3) [10], acquired from institutions all over the world, related to all aspects of the disease, including risk assessment of being infected, triage at hospital admission, prediction of recovery process during follow-up, and patient inclusion and stratification in clinical trials. We aim to be inclusive, and thus models that are outside the scope of risk assessment and patient stratification are still within the purview of the platform, e.g., diagnostic models. We believe it will be of interest to doctors who want to leverage the results of all the great research that is taking place, and it will also benefit researchers in dissemination of their own work and in learning about the findings of other groups.

The proof-of-concept of such a platform forms the basis of this paper. We intend to maintain this platform as a public service, and increase the number of curated models by encouraging other researchers to share their work through our platform. The benefits to them include (a) helping the researchers to generalize their models by allowing the models to be tested by research groups that are different from the ones that created the model (TRIPOD 4), and (b) improved visibility of their model, which should stimulate usage and citations[11].

## Methods

We reviewed the PubMed database of the National Center for Biotechnology Information (NCBI) and the World Health Organization (WHO) database for COVID-19 publications from December 2019 to June 2020. To find relevant publications to our focus we used the terms in the search field: “COVID 2019 prognostic models”, “novel coronavirus 2019 diagnostic tools”, “COVID-19 predictive models”, and “machine-learning COVID 19 models”.

The steps that we followed from the literature search until the final stage of publishing online are shown in Figure 1.

**Figure 1.**
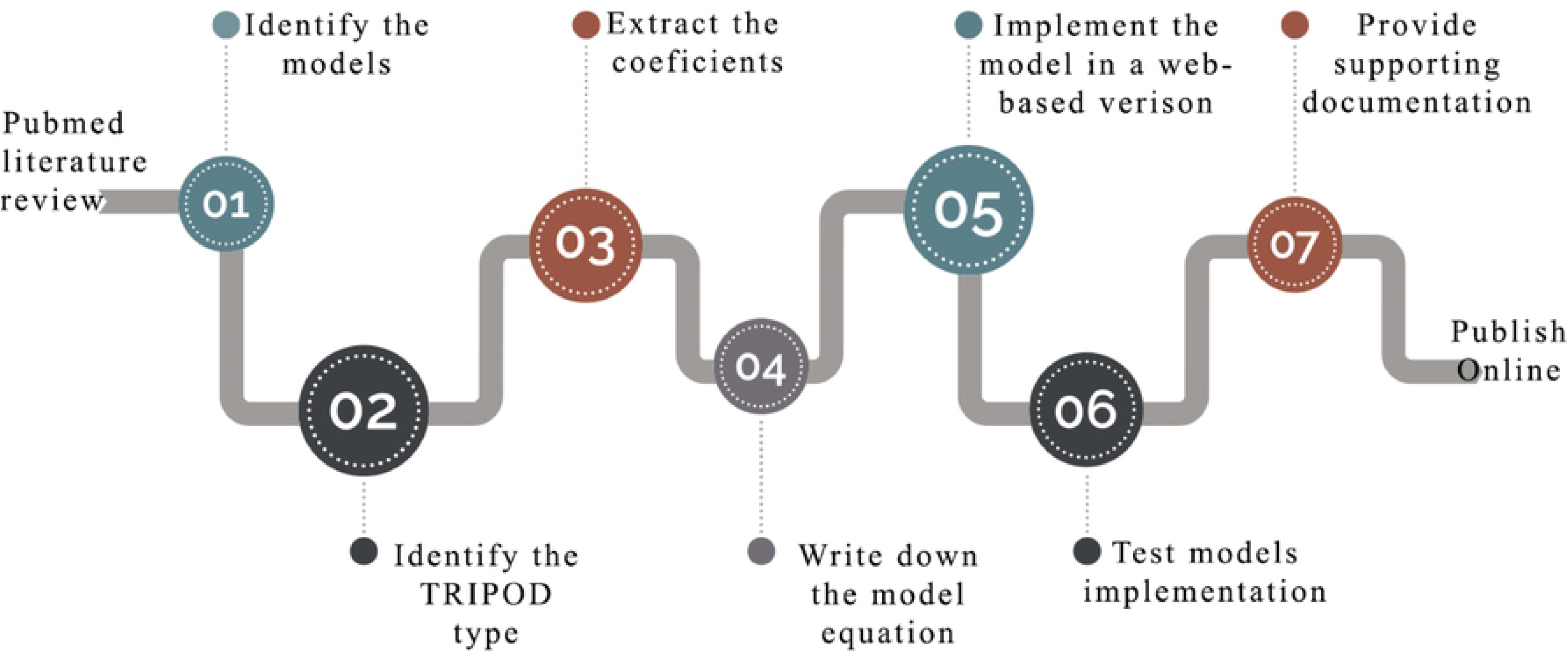
The workflow from defining the convenient models until the end phase.

In order to assess the reporting quality of the models from the studies, we tested each paper for its compliance to the TRIPOD (Transparent Reporting of studies on prediction models for Individual Prognosis Or Diagnosis) reporting guideline as shown in Figure 2 [10,12].

**Figure 2.**
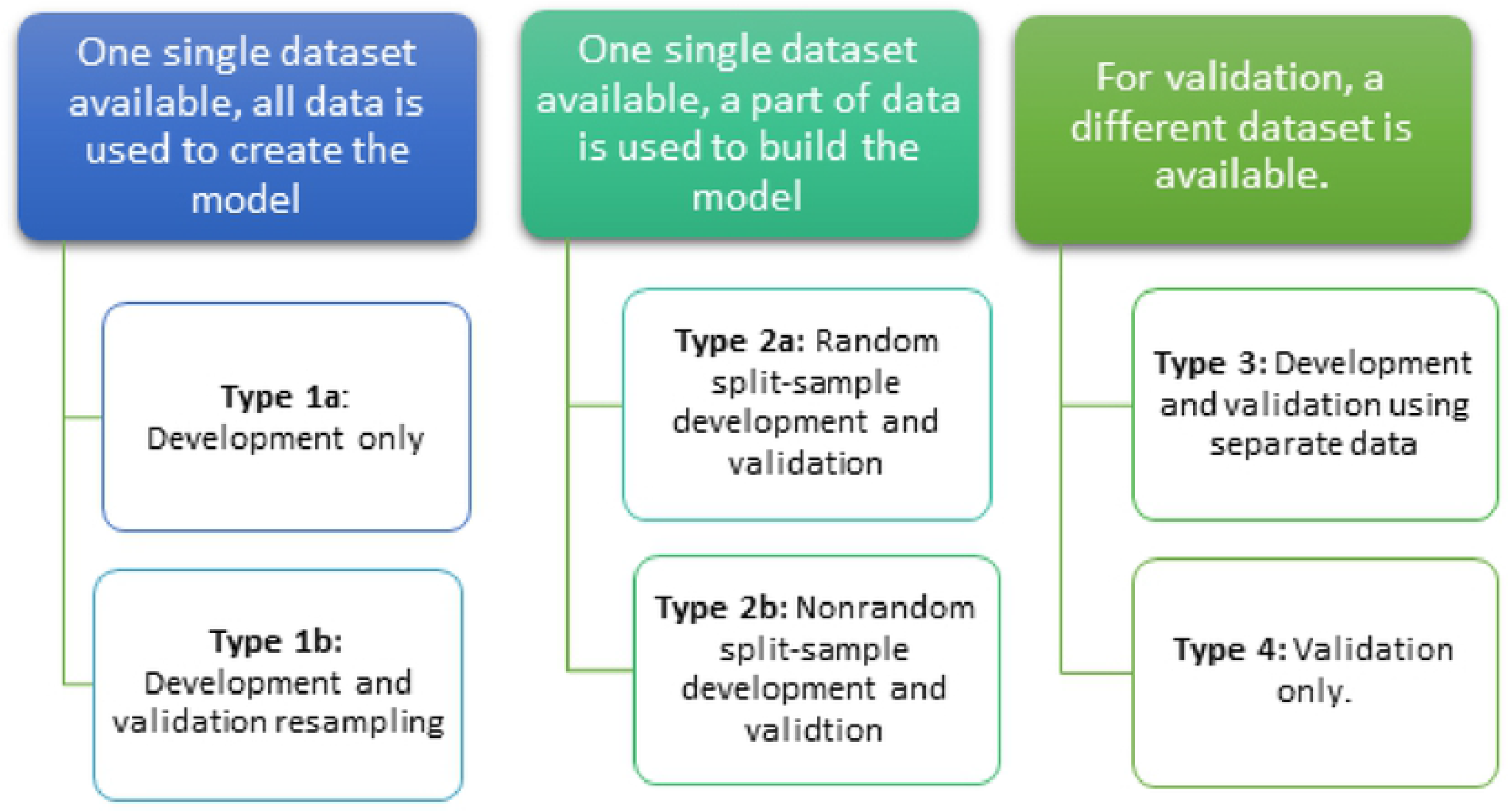
TRIPOD types classifications [10].

### Getting model coefficients from a Nomogram

In order to improve readability and interpretability by medical specialists, regression models are often published as nomograms, without the model coefficients. To publish the models in a consistent manner on our platform, we used a simple method to extract the coefficients from nomograms. This method is explained using an example taken from one of the implemented models [3], and shown below in Figure 3.

**Figure 3.**
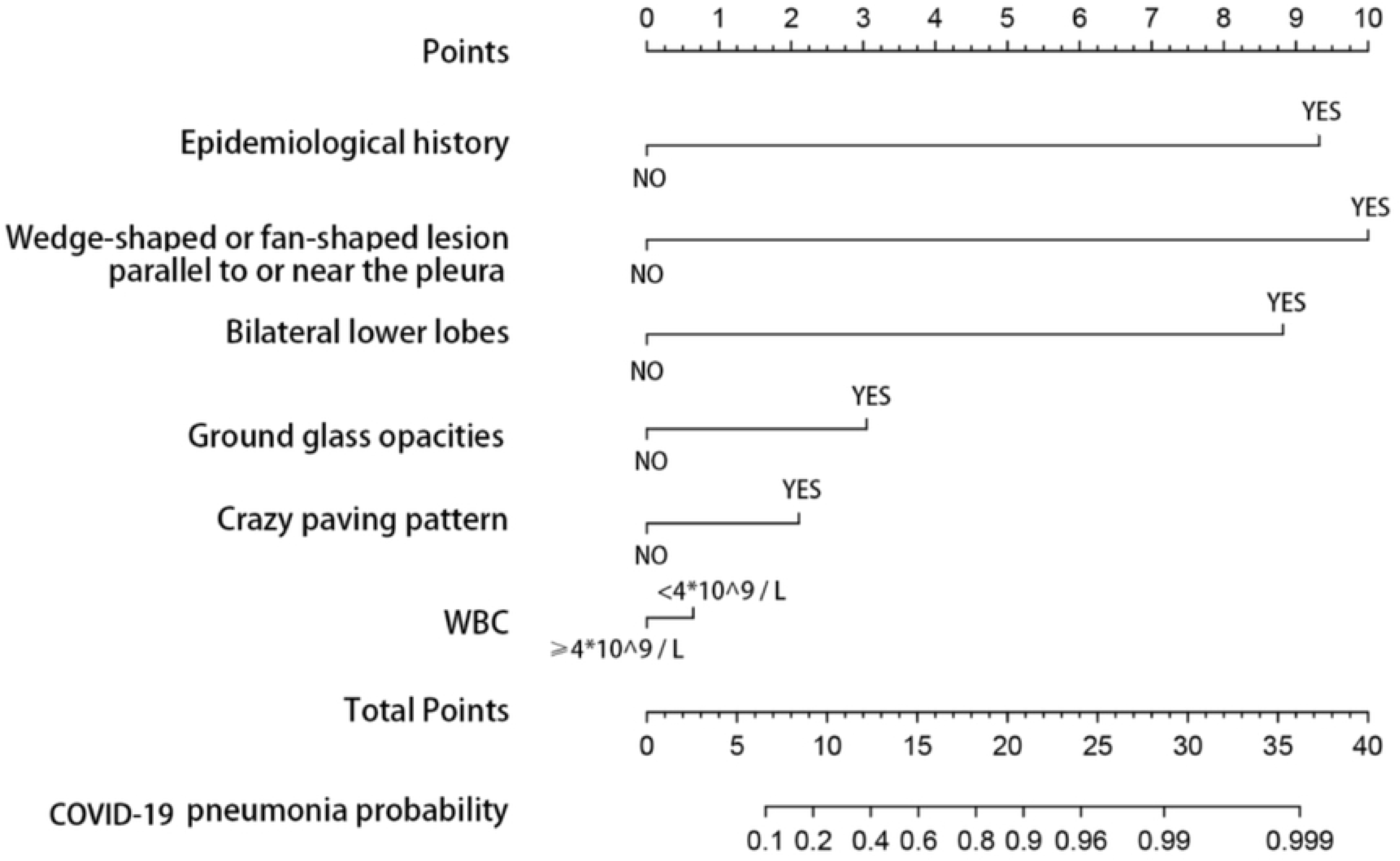
Nomogram Published in https://doi.org/10.1101/2020.04.03.20052068

The first step was to determine the relationship between the parameter and the nomogram score, which was done by reading the nomogram, as shown in Table 1.

**Table 1.** Point reading for the nomogram

| Parameter (Unit) | Equation | Value | Points |
| --- | --- | --- | --- |
| *Epidemiological history*<br>(yes/no, Boolean) | $x_1$ | 1 | 9.32 |
|  |  | 0 | 0.00 |
| *Wedge/fan-shaped lesion*<br>(yes/no, Boolean) | $x_2$ | 1 | 10.00 |
|  |  | 0 | 0 |
| *Bilateral lower lobes*<br>(yes/no, Boolean) | $x_3$ | 1 | 8.82 |
|  |  | 0 | 0.00 |
| *Ground glass opacities*<br>(yes/no, Boolean) | $x_4$ | 1 | 3.04 |
|  |  | 0 | 0.00 |
| *Crazy paving pattern*<br>(yes/no, Boolean) | $x_5$ | 1 | 2.10 |
|  |  | 0 | 0.00 |
| *WBC*<br>($<4 \cdot 10^9/L$, Boolean) | $x_6$ | 1 | 0.63 |
|  |  | 0 | 0.00 |

The relationship between the parameters and the nomogram score *P*_*total*_ is described by the following equation:

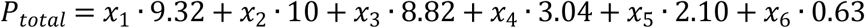

The next step is to determine the relationship between the nomogram score and the probability through the regression equation. A logistic regression model follows the following equation:

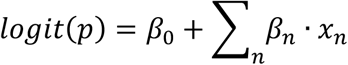

The Logit of the probability and the nomogram score should have a linear relationship, from which the slope was used to determine the value of the coefficients, and the intercept of the model was extracted (Figure 4).

**Figure 4.**
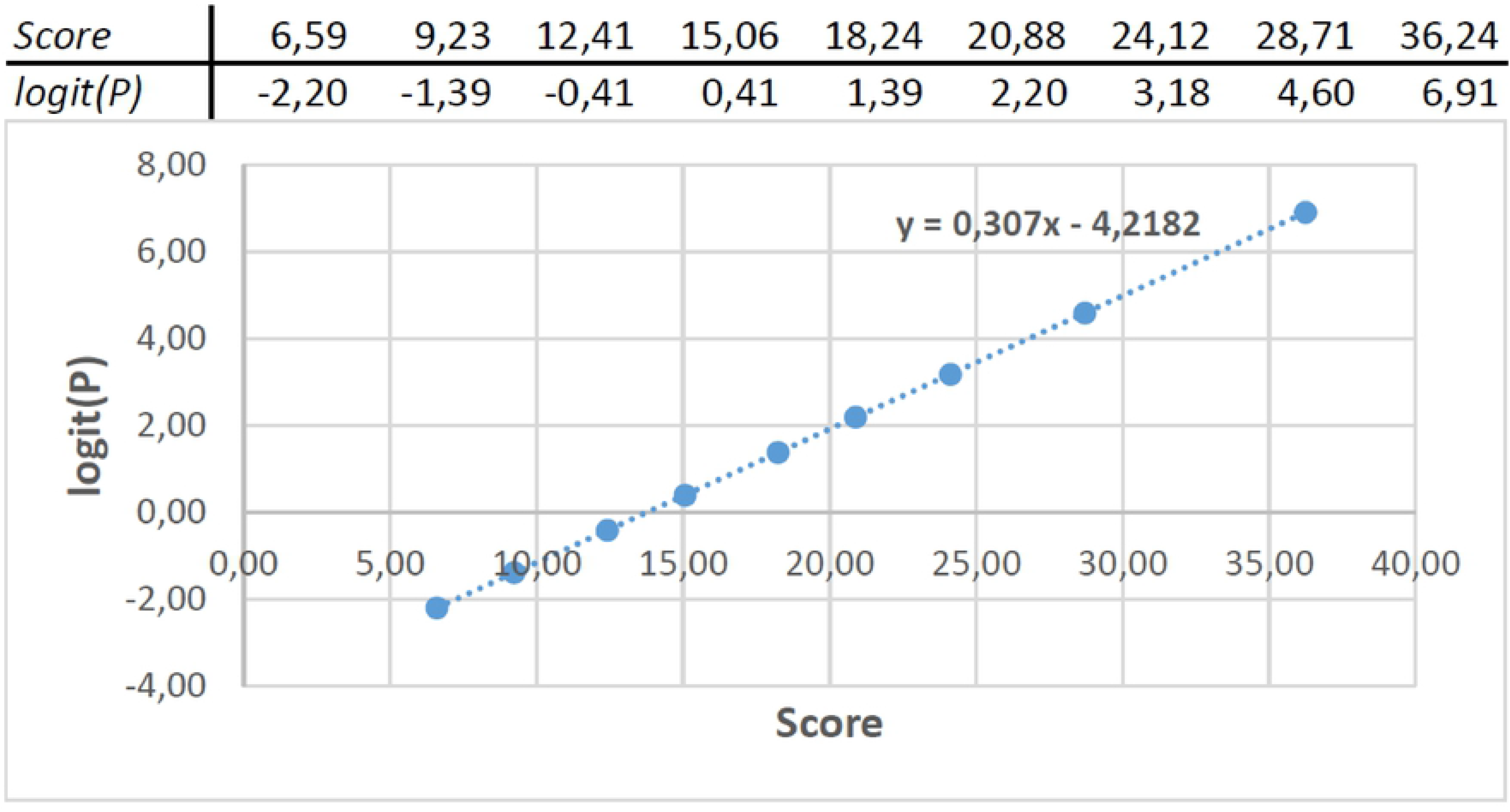
Logit(P) plotted against the nomogram score

For this example the regression coefficients are shown in Table 2.

**Table 2.** Coefficients and intercept extracted from nomogram.

| Parameter (Unit) | Coefficient |
| --- | --- |
| *Epidemiological history* | 0,93 |
| *Wedge/fan-shaped lesion* | 0,64 |
| *Bilateral lower lobes* | 0,19 |
| *Ground glass opacities* | 0,93 |
| *Crazy paving pattern* | 0,64 |
| *WBC* | 0,19 |
| *Intercept* | -4,23 |

All the models are written in PHP 8.0.0, where for regression models we set the coefficients and variables in the PHP syntax, thereby making the models operate identically. For the frontend side we used languages such as HTML, CSS and JavaScript for some specific functionalities. The backend of this platform is PHP based and the database is MySQL.

## Results

We have created an open source website (https://covid19risk.ai/) to serve as an archive for published AI prediction models related to all aspects of COVID-19, including diagnosis, theragnosis (how to treat the patient, risk stratification), and follow-up (treatment response and complication).

Currently there are nine models implemented and published as illustrated in Table 3. Every showcased model includes a description of the methodology and clinical datasets used for model development and validation, and limitations of each model are explicit.

**Table 3.**
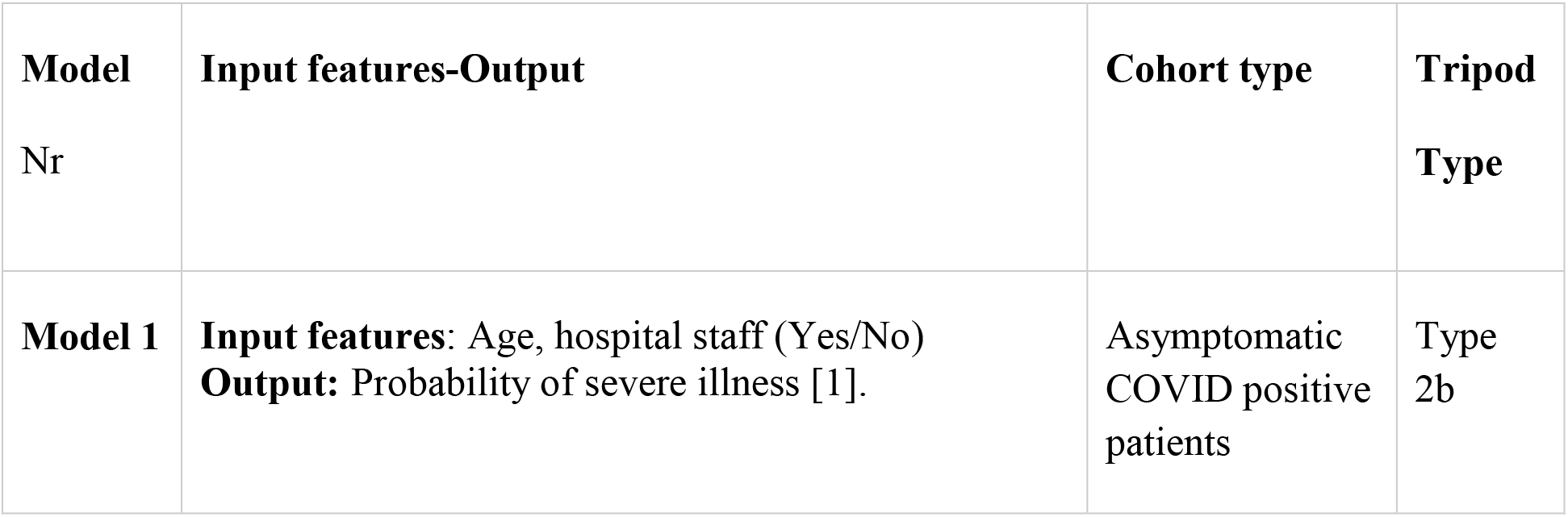
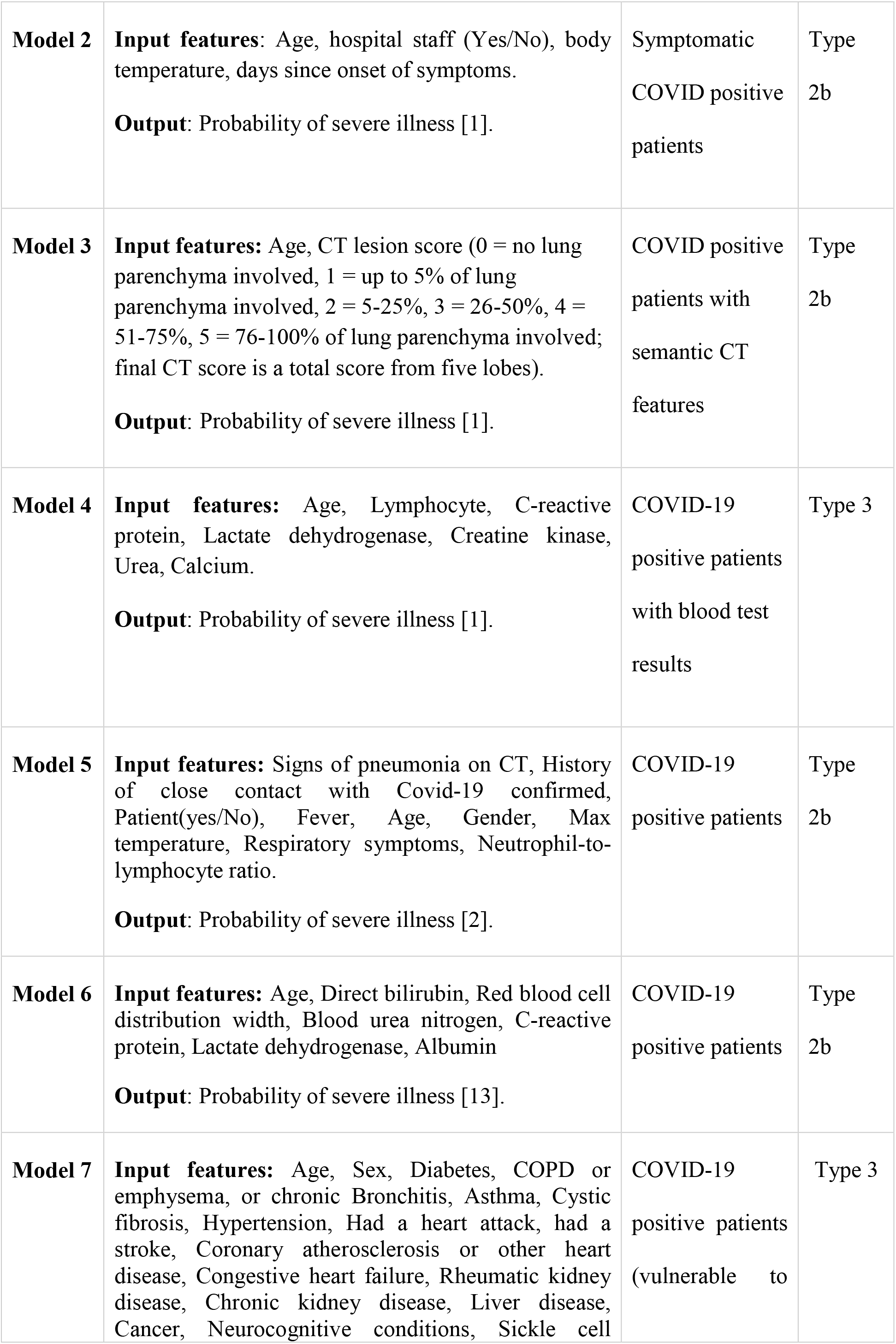
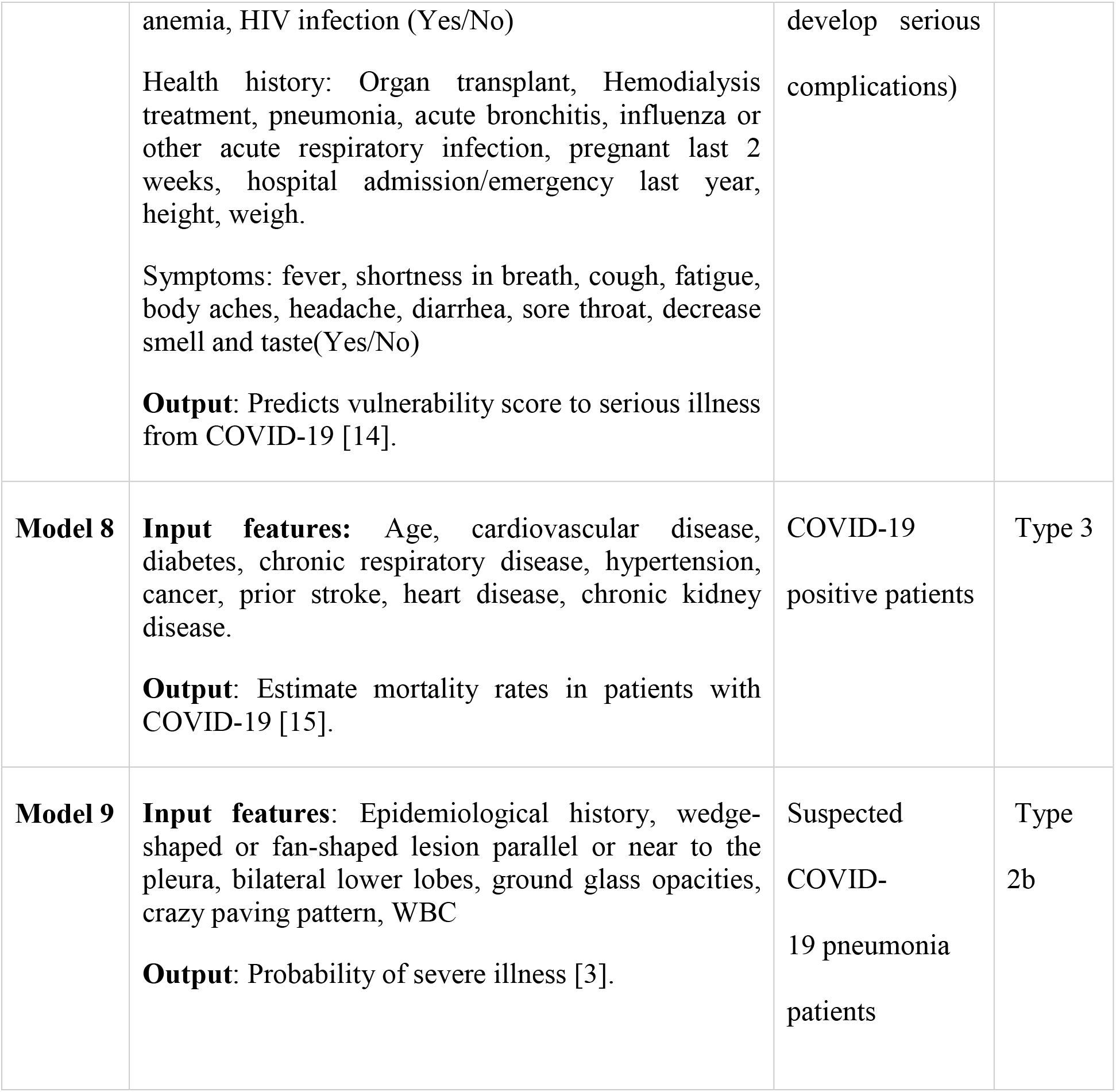
For every model: input features, output, cohort type and TRIPOD type.

For each online model given, doctors can find: a) the intended use (predicted outcome) of the model, b) to which patients does this tool apply (particularly among individuals with preexisting medical conditions), c) the information and the parameters that need to be entered by the doctor, and d) how the tool was developed. The doctors can visit the website, choose an applicable model, and fill in the variables asked in order to generate a probability. The COVID-19 predictive models on the website use the same calculations as the models described in the scientific publications on which they are based.

The main result of our work is a broadly applicable platform, which includes validated models regarding different stages, symptoms and outcomes of COVID-19. This repository of COVID-19 predictive models will serve as a decision aid for doctors.

## Discussion

This platform can be viewed as a “model zoo” aimed at researchers and clinicians and with adequate grasp of the medical complexities associated with COVID-19. The aim for all showcased models is to stimulate research and supplement clinical judgment, not substitute it. The open source website is not intended for unaided use by laypeople (e.g., patients).

We re-emphasize that this manuscript and the website in its current form are only a proof-of-principle. We do not claim that all models that would pass our selection criteria have been included. Similarly, any model not currently included on the platform should not be seen as problematic. Our inclusion period ranged from December 2019 till June 2020. As many models were published since then, an update of the search and the website needs to be and will be done in the near future.

This paper should be seen by researchers from outside our collaboration as an invitation to participate on this platform, with the option of keeping the code hidden from the end user while still offering full functionality. We will assist external researchers for the successful incorporation of their models on our platform. This will create synergies that are bound to accelerate AI research on COVID-19. It will also ensure that models get the recognition they deserve and are used widely, instead of gathering dust as often happens when there are many publications on the same broad theme during a short period (a certainty in the context of COVID-19, given its world-changing nature).

The method we used for retrieving coefficients of a regression model from a nomogram has certain limitations. For one, the accuracy is highly dependent on the resolution of the published model. Another limitation is that though the coefficients of the model are retrieved, the standard error for the coefficients of the parameters cannot be obtained from a nomogram alone. However, the method can be applied to any nomogram, making it a tool that can be broadly used, not restricted to COVID-19.

## Conclusions

Our platform (https://covid19risk.ai/), at the current proof-of-principle stage, includes nine validated machine-learning models to serve as decision aids to doctors for various aspects of COVID-19 patient care. Our method for obtaining regression coefficients from a nomogram can be used by other researchers, including in non-COVID contexts. Our platform will be maintained and regularly updated for at least three years, since we have secured funding for this period (DRAGON grant). Therefore, we are encouraging research groups to collaborate with us to share their models with the world.

## Acknowledgments

Authors acknowledge financial support from the European Commission’s Horizon 2020 research and innovation programme under grant agreement MSCA-ITN-PREDICT n° 766276, and the IMI2 Joint undertaking program under grant agreement DRAGON n° 101005122.

